# Analysing the Role of KIN10 in Directional Root Growth Regulation in *Arabidopsis thaliana*

**DOI:** 10.1101/2025.04.25.650732

**Authors:** Katarzyna Retzer, Judith García-González, Karel Müller, Jozef Lacek, Simon Pree, Nathalie Friedl, Wolfram Weckwerth

## Abstract

Root growth directionality is critical for plant survival, optimizing anchorage and resource acquisition. While the role of hormonal signaling in root gravitropism is well established, the contribution of metabolic status remains less understood. Here, we investigate the function of the catalytic SnRK1 subunit KIN10 in integrating carbon availability with root growth regulation in *Arabidopsis thaliana*. A combination of growth phenotyping, transcriptomics, and hormonomic profiling suggest that KIN10 loss disrupts energy-linked developmental processes. Compared to wild-type *Col-0, kin10* displayed reduced sensitivity to glucose-induced root growth inhibition. Transcriptomic analysis of *kin10* roots revealed widespread reprogramming of metabolic and hormonal pathways, with significant changes in secondary metabolism, cell wall remodeling, and hormone-related gene expression. Hormone profiling further indicated that KIN10 modulates auxin and jasmonate pathways in a carbon source- and organ-dependent manner, especially under sucrose supplementation. Our results demonstrate that KIN10 plays a central role in integrating energy status with developmental and environmental signaling.

## 1. Introduction

Plants continuously adapt their growth and development to a dynamic environment by integrating external cues such as light, temperature, and gravity with internal metabolic status (1–3). In the root system, this ability is vital for optimizing soil exploration, nutrient uptake, and overall plant fitness (1). Gravitropic responses, whereby roots sense the gravity vector and adjust their growth direction accordingly, are crucial for anchorage and efficient resource acquisition (4–6). Beyond the classical hormonal framework, increasing evidence suggests that metabolic status also modulates root developmental responses (7). Yet, the molecular players coordinating energy sensing with gravity-driven growth decisions remain poorly characterized.

SnRK1 (Sucrose non-fermenting-1-Related Kinase 1) has emerged as a central regulator of energy homeostasis in plants (1,7–10). Acting as a metabolic master switch, SnRK1 integrates environmental and internal signals to orchestrate the balance between anabolic and catabolic processes. Under energy deprivation or stress conditions, SnRK1 activates pathways that conserve resources and promote survival, while repressing energy-consuming growth programs (1,10). Structurally, the SnRK1 complex consists of a catalytic α-subunit and regulatory β- and γ-subunits. In Arabidopsis thaliana, two genes encode the catalytic subunits: SnRK1.1 (also known as KIN10 or SnRK1α1) and SnRK1.2 (KIN11 or SnRK1α2) (1,10). Although partially redundant, KIN10 is considered the major active isoform across many tissues and developmental stages (1). Previous studies have shown that SnRK1 activity is crucial for regulating fundamental developmental processes, including root growth regulation (7). Notably, SnRK1’s localized function in the root epidermis, highlights its ability to couple tissue-specific growth responses with the organism’s overall energy status (7). These findings point to SnRK1, and KIN10 in particular, as key integrators of metabolic information and cell-specific developmental outputs.

In this study, we are investigating the role of KIN10 in linking carbon status to root growth pattern depending on illumination status and sugar supplementation to the growth medium in *Arabidopsis thaliana* roots. To uncover potential molecular mechanisms regulated by the cellular master regulator SnRK1, we additionally integrated transcriptomic and hormonomic analyses.

Understanding how energy metabolism intersects with gravity sensing will shed light on the broader integration of metabolic and developmental networks in plants to advance our knowledge of SnRK1/KIN10 functions in root development.

## 2. Results

### 2.1. KIN10 root growth patterns under sugar supplementation

In a previous study we showed that root growth pattern differ in wilde-type roots depending on illumination status of shoot and root, as well on the type of sugar supplemented to the growth medium (2). Shading cotyledons limits the availability of endogenous photosynthates, leading to suppressed root meristem activity and overall shorter root systems (2). However, exogenous supplementation with carbon sources such as sucrose or glucose can counteract this growth inhibition. Sucrose supplementation was shown to more effectively promote both root and hypocotyl elongation compared to glucose, particularly under conditions of increased shading or complete etiolation. Interestingly, while sucrose consistently enhanced root growth regardless of light conditions, glucose supplementation only improved root length when cotyledon illumination was severely restricted (2). This indicates that not only the presence of sugars but also their specific type critically influences root development. These findings highlighted the complex crosstalk between light signals and sugar availability in regulating root architecture (2).

In this study we compared the respective responses of an established KIN10 knockout mutant (*kin10*) to its corresponding wild-type *Col-0*. In the experimental settings, distinct illumination conditions were used to differentiate sugar releated effects: dark-grown roots (DGR), light-grown roots (LGR), and etiolated seedlings. DGR plants are cultivated with roots shielded from light, promoting enhanced meristematic activity and resulting in longer roots compared to LGR, where direct root illumination suppresses root growth by inhibiting cell proliferation and altering hormonal signaling pathways. Etiolated seedlings, which are grown entirely in darkness, display elongated hypocotyls and severely limited root development due to the prioritization of shoot elongation in search of light.

The experiment testing *kin10* root growth adaptation under the differnet growth conditions compared to *Col-0* responses, showed a tendency of *kin10* to react less sensitive to glucose induced root growth inhibition under DGR and LGR conditions (Fig. 1A). Furthermore, also a tendency to react less intensive in terms of root growth deviation from vertical, as indicated by the gravitropic index (GI), was observed for *kin10* roots, especially under DGR and sugar supplementation (Fig. 1B).

**Figure 1:**
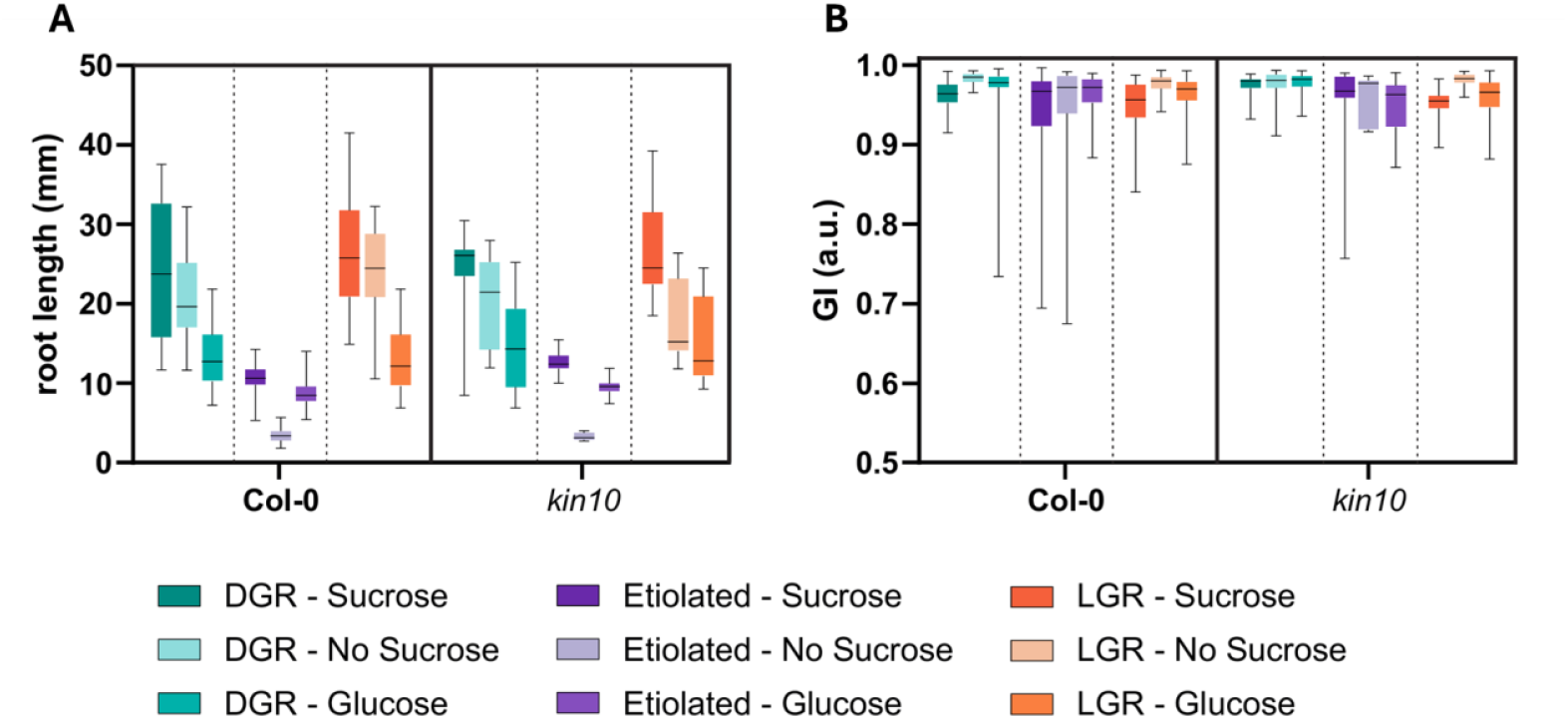
Comparative analysis of root growth responses between wild-type Col-0 and the KIN10 knockout mutant (kin10) under different illumination and sugar supplementation conditions. (A) Total root length of 7DAG *Col-0* and *kin10* seedlings grown under dark-grown root (DGR), light-grown root (LGR), and etiolated conditions, with and without 1% glucose or 1% sucrose supplementation to the MS growth medium. DGR seedlings were cultivated with roots shielded from light, whereas LGR seedlings had roots exposed to direct illumination. Etiolated seedlings, grown in complete darkness, displayed elongated hypocotyls and therefore limited root development. *kin10* mutants showed a reduced sensitivity to glucose-induced root growth inhibition compared to *Col-0* under both DGR and LGR conditions. (B) Gravitropic index (GI) analysis revealed that *kin10* roots showed the tendency for weaker deviation from vertical growth compared to *Col-0*, particularly under DGR conditions with sugar supplementation, indicating altered sugar-dependent regulation of root orientation.

### 2.2. Transcriptomic reprogramming in *kin10* roots reveals metabolic and hormonal disruption

To investigate the molecular mechanisms that underpin root growth differences, we performed RNA-seq analysis of *Arabidopsis thaliana* roots from seedlings grown 7DAG in DGR conditions, on 1% glucose supplemented growth medium. The analysis revealed substantial transcriptional reprogramming in *kin10* roots compared to *Col-0* wild type. The differerently regulated biologcal processes are summarized in Figure 2, highlighting significant shifts in metabolic, hormonal, and stress-response pathways. Upregulated genes in *kin10* roots are associated with secondary metabolism, cell wall remodeling, and stress adaptation. Notably, genes involved in flavonoid biosynthesis and cell wall modification, were induced, suggesting a compensatory shift toward reinforcing cell walls and modulating growth. This needs to be further investgated to understand if this correlates to the alterd growth pattern of *kin10* roots. Downregulated genes in *kin10* roots included several involved in hormone homeostasis, which may contribute to impaired auxin gradients essential for gravitropic bending. This should be also further invastigated in follow-up studies.

**Figure 2:**
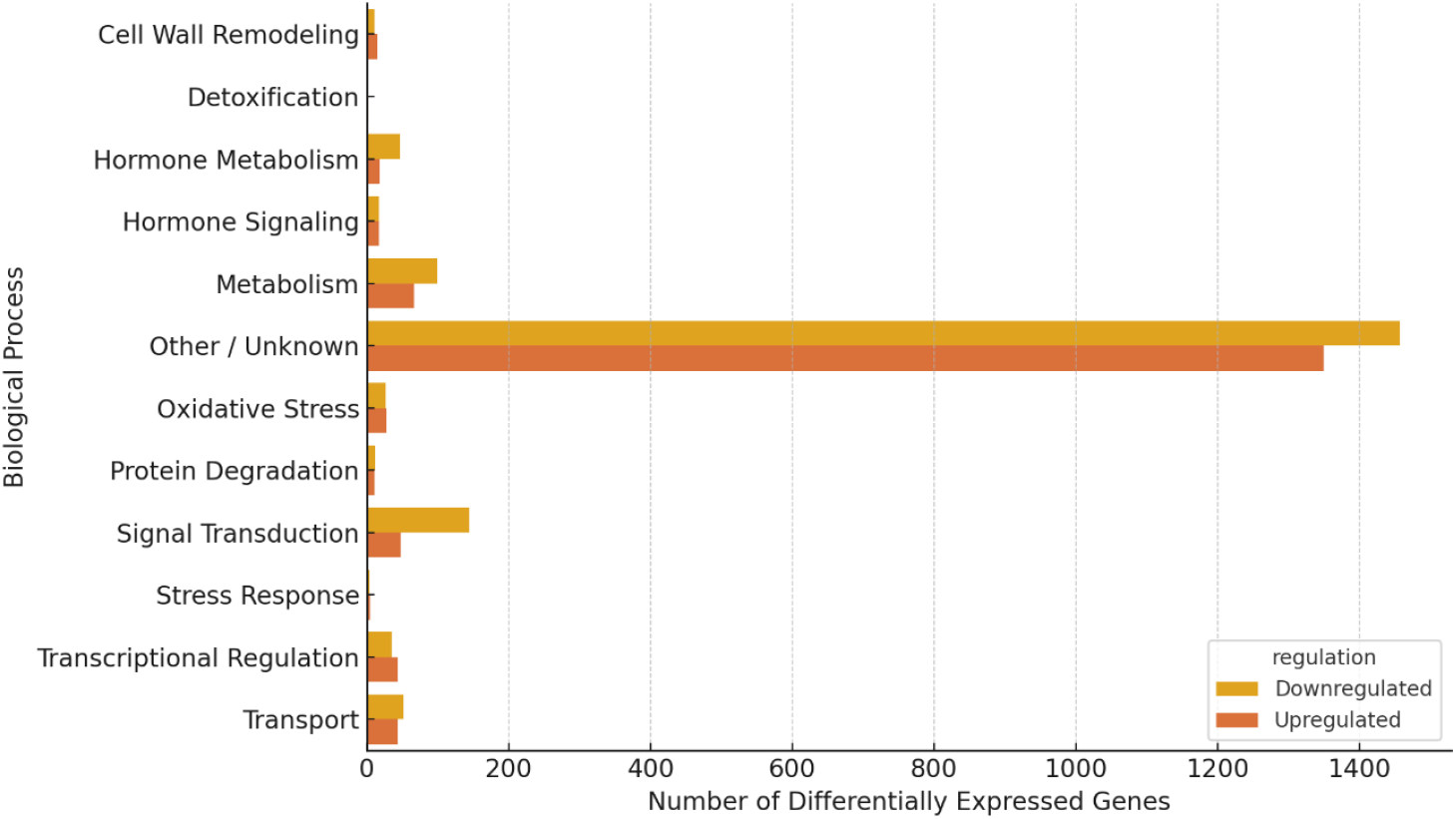
Differential gene expression in kin10 roots compared to Col-0 reveals transcriptional rewiring. Bar plot showing the number of upregulated (orange) and downregulated (yellow) genes in each major biological process category, based on RNA-seq analysis of 7DAG *Arabidopsis thaliana kin10* roots compared to *Col-0*. Genes were classified into categories based on functional keywords in their annotations. While a large fraction of genes fell into the “Other / Unknown” category, many differentially expressed genes clustered into defined processes including metabolism, hormone metabolism and signalling, transcriptional regulation, oxidative stress response, and cell wall remodelling. These results reflect widespread transcriptional reprogramming in kin10 roots, consistent with a central role for KIN10 in coordinating energy signalling with developmental and stress-related pathways.

The transcriptomics dataset indicates that SnRK1 is essential for maintaining transcriptional homeostasis in the root, and further studies are requires to test ist function in integrating energy status with hormonal signaling and developmental control to underpin driectional root growth control.

### 2.3. KIN10 modulates hormone levels and metabolism in an organ- and sugar-dependent manner

To investigate whether loss of KIN10 influences hormone homeostasis as indicated by the RNAseq experiment, we profiled a broad spectrum of hormone metabolites in shoots and roots of 7 DAG *Arabidopsis thaliana Col-0* and *kin10* seedlings grown under different carbon supply conditions. Plants were cultivated on MS medium without sugar, or supplemented with 1% glucose or 1% sucrose, and hormone levels were quantified. As shown in Figure 3, roots exhibited the most pronounced differences between *kin10* and wild-type plants, particularly under sucrose supplementation (Fig. 3B). *kin10* roots accumulated significantly higher levels of compounds belonging to the JA pathway, suggesting a strong KIN10-dependent regulation of these pathways under elevated carbon supply.

**Figure 3.**
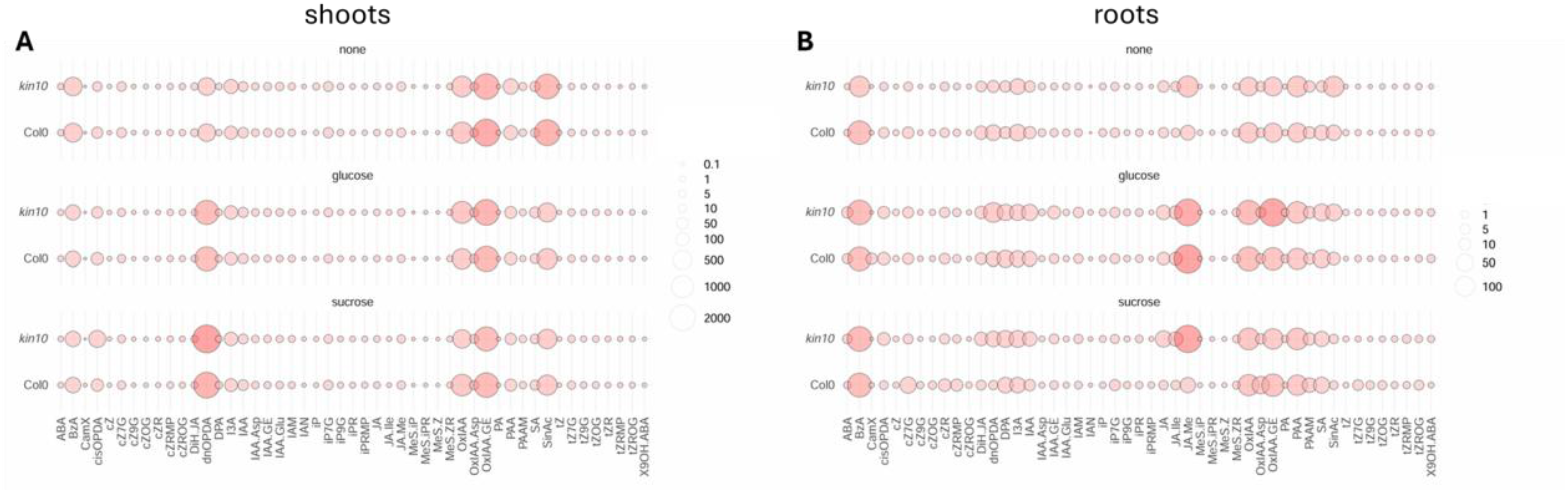
KIN10 modulates hormone abundance in a sugar- and tissue-dependent manner. Bubble plots show relative hormone expression values measured by LC-MS/MS in (A) shoots and (B) roots of *Arabidopsis thaliana Col-0* (wild-type) and *kin10* 7 DAG seedlings. Plants were grown on MS medium without sugar (top), with 1% glucose (middle), or 1% sucrose (bottom). Each bubble represents a specific hormone metabolite, with size and color intensity corresponding to the abundance (log scale). In roots (B), *kin10* shows stronger divergence from wild-type under sucrose supplementation, particularly for auxin- and jasmonate-related compounds. These data highlight KIN10’s role in modulating hormone levels based on organ and carbon source availability.

These findings support a model in which KIN10 differentially modulates hormonal pathways in an organ-specific and carbon-source-dependent manner, aligning with its known role as an integrator of energy status and environmental responses.

## 3. Discussion

Root system architecture is critical for plant survival, enabling efficient resource acquisition and anchorage. The regulation of directional root growth is a highly integrated process, classically attributed to hormonal gradients, especially auxin. However, our study sheds light on a new regulatory layer, demonstrating that energy status, via KIN10, a catalytic subunit of the SnRK1 complex, substantially modulates root growth directionality in Arabidopsis thaliana. Integrating phenotypic, transcriptomic, and hormonomic analyses, we suggest that KIN10 acts as a crucial mediator linking carbon availability to developmental responses.

Our transcriptomic analysis of *kin10* roots uncovered widespread reprogramming of metabolic pathways, including significant upregulation of genes involved in secondary metabolism and cell wall remodelling. These findings suggest that KIN10 normally acts to restrain energy-intensive biosynthetic pathways under conditions of limited carbon supply. In the absence of KIN10, roots might shift toward reinforcing their structure, potentially to compensate for impaired growth regulation, and further investigations would be required to dissect the role of the SnRK1 complex. Upregulation of flavonoid biosynthesis, for instance, may alter auxin transport and distribution, affecting root growth orientation. Similarly, cell wall remodelling genes, often associated with changes in cell elongation and mechanical properties, could contribute to the observed deviations in gravitropic response. Thus, metabolic rewiring in *kin10* roots likely reflects an adaptive strategy to maintain root function despite altered energy sensing, yet at the cost of precise directional growth.

The profound impact of KIN10 on hormonal pathways depending on sugar supplementation, particularly auxin and jasmonate metabolism, underscores the complexity of the integrative role of SnRK1. Hormonomics revealed organ- and sugar-specific changes in auxin- and jasmonate-related metabolites in kin10 seedlings. Notably, in roots, sucrose supplementation amplified these hormonal discrepancies. These findings suggest that KIN10 fine-tunes the balance between growth-promoting and stress-responsive hormonal pathways, ensuring that energy resources are channelled toward appropriate developmental outcomes.

Our findings position SnRK1 through KIN10 activity as a central node integrating energy sensing with developmental control in root growth modulation. Collectively, our data support a model wherein KIN10 acts as an integrator of carbon status, metabolic reprogramming, and hormonal signalling to direct root growth decisions. Understanding how energy status is translated into developmental decisions is critical for improving plant resilience and resource use efficiency. Future studies should focus on dissecting the precise downstream effectors of KIN10 in root tissues, including targeted analysis of auxin transporters and jasmonate-responsive transcription factors. Moreover, cell-specific studies could reveal how KIN10-mediated signalling integrates across diverse cell layers to achieve coordinated root bending and growth orientation.

## 4. Materials and methods

### 4.1. Plant material and cultivation

Experiments were performed on *Arabidopsis thaliana Col-0* and *kin10* (*Col-0* background). Seeds were surface sterilized in a 30 % bleach solution (SAVO©, Unilever, Czech Repulic) with a droplet of Tween 20 for 5 min, rinsed 4 times with distilled water. For the root experiments, seeds were placed on medium in Petri plates filled with half-strength Murashige-Skoog medium supplemented with 1% agar, 25 seeds per plate. Petri plates were placed vertically in the D-ROOT system (11) to allow root elongation in the dark and cultivated for 7 days at 22°C, 16 h light/8 h dark regime. Plates were scanned to obtain images of the seed-lings, and changes in growth characteristics were quantified using Fiji/Image J (12).

### 4.2. Transcription networks establishment (RNAseq)

Total RNA was isolated from approximately 50 mg of root tissues using RNeasy Plant Mini kit (Qiagen) and treated with DNA-Free kit (Thermo Fischer Scientific). RNA purity, concentration and integrity were evaluated on 0.8% agarose gels (v/w) and by the RNA Nano 6000 Assay Kit using Bioanalyzer instrument (Agilent Technologies).

#### RNA-Seq Workflow

For RNA-seq analysis, approximately 5 µg of RNA were submitted for the service procedure provided by Eurofins Genomics. The analysis resulted in at least 10 million 150 bps read pairs. Rough reads were quality-filtered using Rcorrector and Trim Galore scripts (13).Transcript abundances (transcripts per million – TPM) were determined using Salmon (14) with parameters --posBias, --seqBias, --gcBias, --numBootstraps 30. Index was built from TAIR10 *Arabidopsis thaliana* cds dataset. Visualization, quality control of data analysis and determination of differentially expressed genes were determined using sleuth (version 0.29.0) package in R (15). Transcripts with q-value <= 0.05 and log2 fold change >=1 (upregulated) or <= -1 (down-regulated) were considered to be significantly differentially expressed.

### 4.3. Hormonal analysis

7 DAG seedlings were harvested from ½ MS medium without, or supplemented with 1% sucrose or glucose respectively. Shoots and roots were separated from each other and hormone profiles obtained according to Schmidt et al., 2024 (16).

